# Behaviorally relevant cell ensembles in rat motor cortex are replayed during sleep and implicate hippocampal involvement in motor skill learning

**DOI:** 10.1101/2025.10.17.683181

**Authors:** Peyman Nazari-Robati, Michael Eckert, Masami Tatsuno

## Abstract

Motor memory is essential for our daily activities. It involves complex neural processes during learning and sleep. However, unlike explicit memory where its neural activation and role in memory consolidation are well-studied, the properties of cell ensembles for motor memory are less understood. In this study, we re-examined rats’ behavior and neural activity in the primary motor cortex (M1) and hippocampus while the animals were trained daily on a single-pellet reaching task. Recordings included both the training and 3 hr rest epochs before and after training. Behaviorally, the animals were classified into two learning types: rapid and gradual learners. Unsupervised cell ensemble detection on M1 neurons revealed that about 60% of the ensembles were modulated during reaching behavior. Those reach-related ensembles were further categorized into four types, and their replay was detected during both slow-wave sleep (SWS) and REM sleep. In SWS, replay preferentially occurred during spindles, especially slow-oscillation coupled spindles (SO-spindles). In addition, about 30% of the reach-related cell ensembles were modulated during the hippocampal sharp-wave ripples (SWRs). The direction of modulation and the temporal coupling between SWRs and SO-spindles depended on the training phase and the animals’ learning types. Our results demonstrate the replay of rats’ skilled-reaching memory during SWS and REM sleep and the possible involvement of the hippocampus through the modulation of M1 activations during SWRs. This study will advance our understanding of how neural activity patterns evolve during skilled-reaching learning and sleep, and help develop medical applications that leverage sleep’s memory functions.

**Significance Statement:** Most evidence for memory replay comes from hippocampal studies in which neural activity during rest is compared to a template constructed from activity recorded during a behavioral task. Here, we used an unsupervised ensemble detection on recordings from rat primary motor cortex (M1) that included both rest and a skilled reaching task. We discovered a variety of ensembles with different sizes and time scales. Many were related to reaching behavior and were replayed during sleep. Some M1 ensembles were also modulated by the hippocampal activity, suggesting their involvement in motor skill learning. This study will advance our understanding of motor memory and sleep and help develop medical applications that leverage sleep’s memory functions.

## Introduction

Memory is a vital brain function for both humans and animals. Our daily activities depend heavily on memory, which serves as the foundation for higher-order brain functions. Memory is typically divided into explicit memory, which involves the conscious recall of facts and events, and implicit memory, which involves the unconscious recall of skills and motor behavior (Squire, 2004).

The conjecture that memory reactivates during rest is supported by findings that behaviorally induced neural activity reactivates during quiet wakefulness and sleep (Wilson and McNaughton, 1994; Foster and Wilson, 2006). Here, we use reactivation as a general term that describes the re-emergence of prior neural activity at a later time point, and replay as a specific form of reactivation characterized by sequential activity (Genzel et al., 2020).

Reactivation of explicit memory has been extensively studied. It has been observed in various brain regions, including the hippocampus and neocortex, during quiet wakefulness and slow-wave sleep (SWS) (Wilson and McNaughton, 1994; Foster and Wilson, 2006), particularly during sharp-wave ripples (SWRs) in the hippocampus and spindles in the cortex (Ji and Wilson, 2007; Peyrache et al., 2009). Disruption of reactivation results in impaired memory consolidation, supporting the hypothesis that coordinated reactivation in the hippocampus and the neocortex plays an important role in the consolidation of explicit memory (Girardeau et al., 2009; Ego-Stengel and Wilson, 2010; Gridchyn et al., 2020).

For implicit memory, several studies have reported reactivation of motor skill memory in the primary motor cortex during SWS (Ramanathan et al., 2015; Eckert et al., 2020) as well as REM sleep (Eckert et al., 2020). Reactivation was often observed during spindles (Ramanathan et al., 2015; Eckert et al., 2020), and the density of slow-oscillation coupled spindles (SO-spindles) increased after motor learning (Kam et al., 2019). In these studies, reactivation was detected using principal component analysis (PCA). However, since PCA is not sensitive to sequential patterns, it remains unclear whether the sequential neural activity observed during motor behavior is replayed during rest epochs. In our previous study (Eckert et al., 2020), we attempted to detect replay using template matching (Tatsuno et al., 2006). However, unlike in the case of explicit memory, detection was not successful even though robust sequential activity patterns were observed in M1 during behavior. Therefore, our understanding of the replay of implicit memory is still limited.

Motor learning is a dynamic process that involves several steps, including initial acquisition often marked by rapid performance improvement, and further acquisition and consolidation characterized by the gradual automation of acquired skills (Karni et al., 1998; Hikosaka et al., 2002; Doyon and Benali, 2005). Key brain areas for motor learning include the cerebellum, the basal ganglia, and the primary motor cortex (M1) (Doyon and Benali, 2005). Some research, especially human studies, also suggests the involvement of the hippocampus in motor sequence learning (Albouy et al., 2008; Gheysen et al., 2010; Jacobacci et al., 2020).

Here, we analyzed our previously recorded data (Eckert et al., 2020) to investigate the replay of sequential patterns of cell firing in M1. These are groups of neurons whose activity is significantly correlated during the recording, and they are called cell ensembles. We utilized an unsupervised detection method because it is designed to detect sequential activities without relying on behavioral reference events, unlike template matching. Specifically, we employed an approach proposed by Russo and Durstewitz (2017), which can identify cell ensembles at different time scales, levels of precision, and with arbitrary internal organization. We explored the following questions: What kinds of M1 cell ensembles are detected when rats learn a skilled reaching task and when they rest? Do the reaching-related cell ensembles replay during SWS, REM sleep, and spindles? Is the replay modulated by hippocampal SWRs? If so, are the direction of modulation and the temporal coupling of SWRs and spindles related to how animals learn?

## Materials and Methods

### Animal and Surgery

We analyzed the data previously acquired in our lab (Eckert et al., 2020). Briefly, four adult male Fisher-Brown Norway rats were trained on the single pellet reaching task (Whishaw and Pellis, 1990). Each daily recording session consisted of a 3-hour rest epoch before the task (rest 1), followed by 30 minutes of skill training on the reaching task, and concluded with a 3-hour rest epoch after the task (rest 2) (Figure 1A). A hyperdrive (Gothard et al., 1996) with 12 tetrodes and two reference electrodes was implanted in the forelimb area of the primary motor cortex (coordinates: 1.00 mm anterior, 2.5 mm lateral to bregma), and the reference electrodes were positioned in the cortical white matter for tetrode recordings. Neuronal spike activity was recorded throughout the entire 6.5-hour recording session. In addition, hippocampal local field potential (LFP) was recorded using a twisted bipolar electrode implanted in the dorsal hippocampus. EMG activity was measured with a wire implanted into the neck muscle.

**Figure 1.**
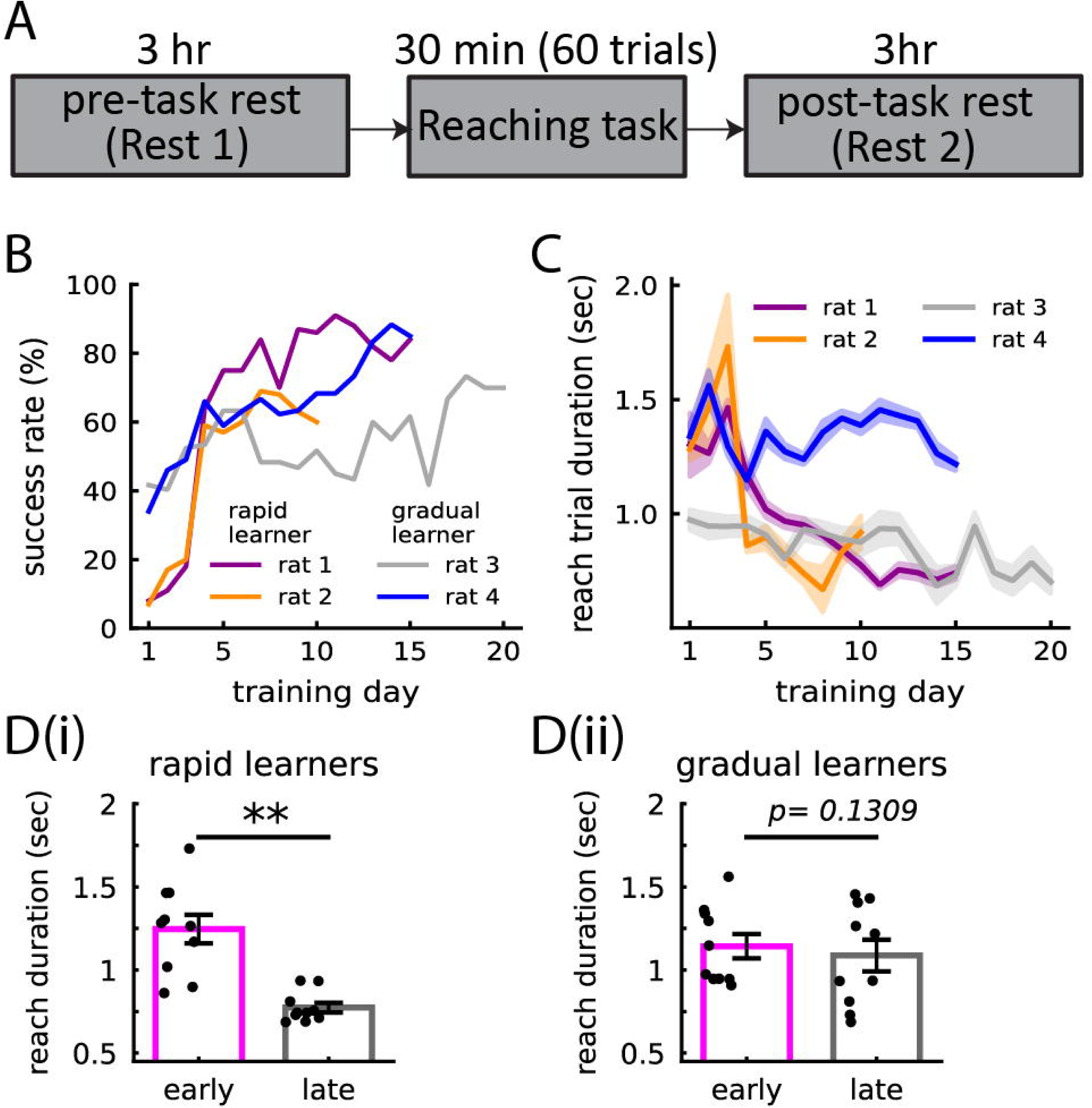
Experimental design and behavioral performance. **A.** Experimental procedure. A recording session consisted of a 3-hour pre-task epoch (rest 1), a 30-minute task epoch with training of the reaching task, and a 3-hour post-task epoch (rest 2). Spiking activity in the primary motor cortex was recorded continuously using tetrodes. **B.** Success rate across learning days. Rats 1 and 2 exhibited low success rates initially, followed by a significant increase on day 4 and gradual improvement thereafter. Rats 3 and 4 showed moderate success rates from the early phase of training, with gradual changes across days. Rats 1 and 2 were classified as rapid learners, and rats 3 and 4 were classified as gradual learners (adapted from Eckert et al., 2020, with permission). **C.** Change of the duration of reaching trials, measured as a lag between paw advance (reach onset) and paw return (reach offset) across training days (mean ± SEM across trials). **D.** Average duration of reaching trials during early and late training phases. **D(i)**. Rapid learners (Wilcoxon signed-rank test, p = 0.0039, mean ± sem). **D(ii).** Gradual learners (Wilcoxon signed-rank test, p = 0.1309, mean ± sem).

### Data acquisition

The recording process employed the digital Cheetah SX data acquisition software by Neuralynx (Boseman, Montana, USA). Spike data were collected using bandpass filtering (600–6000 Hz) and sampled at 32 kHz. LFP and EMG signals were filtered (0.1–1000 Hz), sampled at 2 kHz, and referenced to a cerebellar skull screw. Spikes were sorted through automated clustering (Klustakwik, K. Harris), followed by manual refinement (MClust, David Redish). Several features assessed the quality of sorted units: the inter-spike interval histogram (less than 0.2% spikes in a 2 msec refractory period), cross-correlation with other units (no peak in the refractory period), waveform shape with low variance and consistent firing rates/patterns throughout the recording, and the L-ratios (units with L-ratios greater than 0.12 were excluded) (Schmitzer-Torbert et al., 2005). Table 1 summarizes the number of spike-sorted neurons used in the analysis.

**Table 1.** The number of spike-sorted neurons for each animal over the training day.

| Day | Rat 1 | Rat 2 | Rat 3 | Rat 4 |
| --- | --- | --- | --- | --- |
| 1 | 27 | 19 | 50 | 21 |
| 2 | 51 | 19 | 58 | 22 |
| 3 | 57 | 46 | 58 | 41 |
| 4 | 37 | 34 | 84 | 36 |
| 5 | 50 | 28 | 76 | 37 |
| 6 | 81 | 21 | 72 | 21 |
| 7 | 47 | 33 | 61 | 42 |
| 8 | 45 | 33 | 63 | 25 |
| 9 | 43 | 31 | 70 | 14 |
| 10 | 59 | 47 | 49 | 14 |
| 11 | 65 |  | 70 | 24 |
| 12 | 63 |  | 62 | 19 |
| 13 | 67 |  | 74 | 16 |
| 14 | 67 |  | 74 | 19 |
| 15 | 39 |  | 70 | 26 |
| 16 |  |  | 71 |  |
| 17 |  |  | 75 |  |
| 18 |  |  | 52 |  |
| 19 |  |  | 62 |  |
| 20 |  |  | 65 |  |

### Behavioral Task

During the task epoch, rats were housed in a custom-designed polycarbonate enclosure tailored explicitly for the single-pellet reaching task. This container featured a front opening with a 1.5 cm slot, allowing the rats to reach through and retrieve a 45 mg food pellet situated 1.5 cm away on a 3 cm high shelf. Task performance was recorded using a high-speed infrared camera capturing at 200 frames per second. An infrared beam at the front of the cage detected reaching actions. The animals participated in approximately 60 trials during the 30-minute session. During the intervals between trials, a sliding door was used to block access to the slot, regulating both the duration and the number of trials in each session. In rest epochs, rats relaxed inside a flowerpot furnished with towels, while their activity was monitored and recorded using an infrared security camera. The rats had no prior experience or training in the specific skill being tested, except for the initial assessment to determine their paw preference. Before training began, all animals underwent a one-week habituation period in which they followed the daily recording procedure (pre-rest, task, and post-rest). During the task epoch, the door to the shelf remained closed, and food pellets were scattered on the floor so the rat did not reach for them.

### Behavioral Assessment

Reaching behavior was evaluated using two metrics: the success rate and the duration of reach trials. The success rate is the ratio of successful trials to the total number of attempted trials. A successful trial is defined as one in which the rat successfully grabbed the pellet, brought it to its mouth, and consumed it. Other trials, such as misses or instances where the rat grasped the pellet but dropped it during retrieval, were classified as unsuccessful. The duration of the reach behavior is defined as the time difference between the start and end of the reaching movement (paw advancing and paw returning).

### Sleep Classification

Sleep was categorized into REM and SWS during rest epochs by calculating the ratio of hippocampal theta power to hippocampal delta power and neck EMG power. The timestamps marking the beginning and end of REM episodes were identified as the period when this ratio exceeded the mean plus three standard deviations. The remaining rest epoch was classified as either SWS or quiet wake based on the strength of upper delta power (2-6 Hz, greater than 2 standard deviations).

### Spindle Detection

Low-voltage spindles (LVS) and high-voltage spindles (HVS) were detected using the LFP from one of the tetrode channels. For LVS detection, the LFP was filtered between 10-20 Hz and then squared to obtain a power signal. Peaks in the power signal greater than 1.5 SD were adjusted down to 0.75 SD to create a list of start and end timestamps. Gaps in timestamps below 100 ms were merged, and a minimum duration of 200 ms was required. LVS was detected in a similar manner, filtering the LFP between 6-10 Hz first and requiring the peak power to exceed 3 SD. Although both LVS and HVS were detected, we focused on LVS because they were related to memory reactivation, but HVS were not (Johnson et al., 2010).

### Slow Oscillation (SO) Detection

The LFP was filtered in the delta band (1 – 5 Hz), and a time-shifted (35 ms) difference signal was calculated. This time-shifted signal emphasized large amplitude fluctuations on the timescale of the slow oscillation. Peaks that exceeded the threshold (mean + 3 SD) were regarded as significant slow oscillations (SOs). If LVS and SOs occurred within a 500-ms window, the event was considered a SO-coupled spindle.

### Sharp Wave Ripple (SWR) Detection

The hippocampus LFP was bandpass filtered within the frequency range of 100 to 250 Hz. After squaring the signal to obtain the power, the mean and standard deviation of the power signal were calculated. Periods that exceeded the threshold (mean + 2.5 SD) were considered SWRs. Events lasting longer than 150 msec were omitted, and events separated by less than 25 msec were combined into a single event.

### Cell Ensemble Detection

Cell ensembles were detected using the unsupervised detection algorithm (Russo and Durstewitz, 2017). The detection was performed separately for each recording session, utilizing the entire recording data. We employed this method because it facilitates the detection of cell ensembles over different timescales (in our analysis, [3, 5, 10, 25, 35, 50, 65, 75, 90, 100] msec) and allows for various internal organizations without requiring templates. Detection was accomplished through a rapid parametric test statistic that accounts for non-stationarities in neural activity and an agglomerative clustering algorithm. Detected cell ensembles were categorized into five types: Type I – precise lag-0 synchronization with a single spike per neuron; Type II – precise sequential pattern with a single spike per neuron; Type III – precise spike-time pattern with multiple spikes per neuron allowed; Type IV – rate pattern with temporal structure; Type V – simultaneous rate increase. For each cell ensemble, activation was represented by a binary activation vector based on the bin size at which the cell ensemble was detected. A value of 1 was assigned when cell ensemble activation was detected and 0 otherwise. The timing of activation was defined as the bin edge of the first neuron’s firing within a cell ensemble.

### Activation of cell ensembles during reaching

To investigate how cell ensembles were activated during reaching behavior, the activation within a ±5-second window around the moment of food pellet grasping was analyzed. We used a 10-second duration because it was approximately 10 times greater than the reach duration (1 ± 0.019 sec, mean ± SD across animals), ensuring a comprehensive examination of cell ensemble dynamics. Using the binary vector of cell ensemble activation, peri-event time histograms (PETHs) for each cell ensemble were generated and smoothed with LOWESS regression (Cleveland, 1981). To identify the cell ensembles that were highly active during reaching behavior, we constructed a histogram of the maximum PETH values for all cell ensembles and selected those that fell above the upper third quantile. Finally, to identify common PETH patterns in the remaining cell ensembles, K-means clustering with a squared Euclidean distance metric was applied. The optimal number of cluster centers was determined by varying K in the range of [2, 20] and selecting the K value with the maximum Silhouette score. We performed the initial K-means clustering on each recording session from each rat. Then we constructed a histogram of K values, selecting the most frequent K as the common number of clusters for subsequent analyses.

### Activation of cell ensembles during sleep

The activation of reach-related cell ensembles during sleep was assessed by calculating their activation rates for each SWS and REM episode. We estimated the activation rate by dividing the number of activations by the duration of the episodes. This procedure was carried out for every reach-related cell ensemble and for each recording session.

### Activation of cell ensembles during spindles

The activation of reach-related cell ensembles during spindles was assessed in two ways. First, both isolated and SO-coupled spindles were combined and treated as single-category spindle events. The average activation counts within ±1 second of the center of spindle events were calculated. To evaluate cell ensemble activation between spindles, a 2-second period between spindles during SWS was randomly selected, ensuring it did not overlap with the spindles. The average activation counts of cell ensembles during this period were calculated, and this procedure was repeated 1000 times to create a distribution of cell ensemble activation between spindles. Second, cell ensemble activation was assessed separately for isolated and SO-coupled spindles, allowing a comparison of the activation strength during isolated and SO-coupled spindles.

### Activation of cell ensembles during SWRs

The activation of cell ensembles during hippocampal SWRs was examined by taking a ±150 msec window around the center of the SWRs. The number of activations was counted using a bin size of 10 msec. Cell ensemble activation between SWRs was also assessed by randomly selecting a 300-msec segment and counting the activations within that period. This procedure was repeated 1000 times, generating the distribution of cell ensemble activations between SWRs, referred to as the non-SWR distribution. Using this distribution, the SWR-associated cell ensemble activation was classified into three groups based on the following criteria: If cell ensemble activation exceeded the 95th percentile of the non-SWR distribution, the cell ensemble was categorized as SWR- enhanced. If cell ensemble activation fell below the 5th percentile of the non-SWR distribution, the cell ensemble was categorized as SWR-suppressed. Otherwise, the cell ensemble was labeled as having no SWR modulation. To characterize the overall direction of change, whether enhanced or suppressed, we also introduced the modulation index, defined as the ratio of the number of SWR-enhanced ensembles to the sum of SWR-enhanced and SWR-suppressed ensembles.

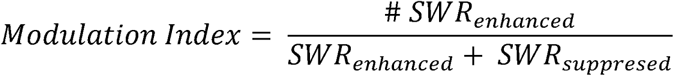

### Replay strength

The replay strength in REM and SWS was assessed using the following measure:

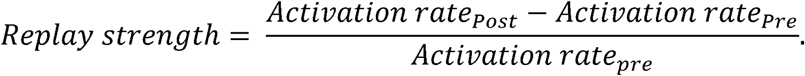

It quantifies the change in cell ensemble activation between the pre-task rest epoch and the post-task rest epoch, normalized by the amount of activation during the pre-task rest epoch.

### Temporal relationship between SWRs and SO-coupled spindles

For each recording session, a peri-event time histogram (PETH) of SWRs was created with respect to the onset of SO-coupled spindles. The midpoint of SWR events was counted using a temporal resolution of 2 msec within a range of ±2 seconds. The PETH was then smoothed with a Gaussian filter with a kernel size of 50 msec. Additionally, the temporal relationship between SWRs and SO-coupled spindles was evaluated by measuring the absolute time difference between the SWR center and the nearest onset of SO-coupled spindles.

### Statistical analysis

The Lilliefors test was employed to assess whether the data followed a normal distribution. If the data did not follow a normal distribution, the Wilcoxon rank sum test was used for comparing independent pairwise samples, the Wilcoxon signed-rank test for comparing repeated pairwise observations, and the Kruskal-Wallis test for comparing more than two repeated groups. Post-hoc multiple comparisons were conducted using Dunn’s test, and p-values were adjusted with the Bonferroni correction. The threshold for statistical significance was set at p < 0.05. Asterisks in the figures indicate the p-value ranges: * 0.01 < p < 0.05, ** 0.001 < p < 0.01, *** p < 0.001. Results were reported as mean ± sem, unless otherwise specified.

### Data analysis software and hardware

The Python version of the cell ensemble detection method (Russo and Durstewitz, 2017), available in the Elephant package—an open-source, community-centered library for analyzing electrophysiological data in the Python programming language, was utilized. Cell ensemble detection was conducted on the Cedar cluster system in the Digital Research Alliance of Canada. The remaining data analysis steps were performed on the Polaris cluster system in the Department of Neuroscience at the University of Lethbridge and on a desktop PC (Intel Core i5-11600K, 32 GB RAM). All programming and data analysis steps were executed using Python 3.10.

## Results

Rats learned the task successfully and were classified into fast or gradual learners First, we investigated how animals’ reaching performance changed over the course of training days. We calculated the success rate of reaching behavior and the speed of the reaching action. All four rats learned the task successfully, as reported in Eckert et al. (2020) (Figure 1B). Two rats exhibited a rapid acquisition of the skill, showing significant performance gains from Day 3 to Day 4, followed by a gradual increase in performance thereafter (Figure 1B, Rats 1 and 2). The other two rats showed a gradual increase in success rate over the entire training days (Figure 1B, Rats 3 and 4). In this paper, we refer to the former as rapid learners and the latter as gradual learners. In addition, the success rate of the rapid learners in the first three days was significantly lower than that of the gradual learners (Wilcoxon rank sum test, p = 0.002).

The speed of the animals’ reaching actions was assessed by measuring the change in duration of the reaching action (Figure 1C). We found that the rapid learners displayed a clear decreasing trend over the training days, while the gradual learners did not show this trend. To quantify these changes, we compared the reaching duration during the early training days (Days 1-5) with the late training days (Days 10-15 for Rats 1, 3, and 4, and Days 6-10 for Rat 2). Statistical tests confirmed that the rapid learners exhibited a significant decrease in duration, but the gradual learners did not (Figure 1D, Wilcoxon signed-rank test, rapid learners (D(i)): n=10, p=0.0039, gradual learners (D(ii)): n=10, p=0.1309).

In summary, all four animals successfully learned the reaching task. The rapid learners showed a significant performance improvement from the initially low success rate within a single day, and their reaching speed decreased significantly between the early and late phases of training. The gradual learners started with an initially intermediate success rate and acquired the task slowly, with their reaching speed remaining steady.

### Unsupervised method detected various cell ensembles

We shifted our focus to the detection of cell ensembles. A total of 60 recording sessions from four animals (rat 1, 15 sessions; rat 2, 10 sessions; rat 3, 20 sessions; rat 4, 15 sessions) were analyzed using the unsupervised cell ensemble detection method (Russo and Durstewitz, 2017). The detection was performed separately for each recording session, utilizing the entire recording data, including pre-task rest, task, and post-task rest epochs. The number of detected cell ensembles in each session was summarized in Table 2. See the supplemental tables for the detected cell ensembles for each rat. Detected cell ensembles were categorized into five types following Russo’s paper: Type I – precise lag-0 synchronization with a single spike per neuron; Type II – precise sequential pattern with a single spike per neuron; Type III – precise spike-time pattern with multiple spikes per neuron allowed; Type IV – rate pattern with temporal structure; Type V – simultaneous rate increase. The overall detection of cell ensembles obtained by summing the detections over different bin sizes was dominated by Type I, II, IV, and V ensembles (Figure 2A). The number of these ensembles was not significantly different, but the number of Type III ensembles was significantly smaller (Kruskal-Wallis test, statistic = 179.44, p=9.83 × 10^-38^).

**Figure 2.**
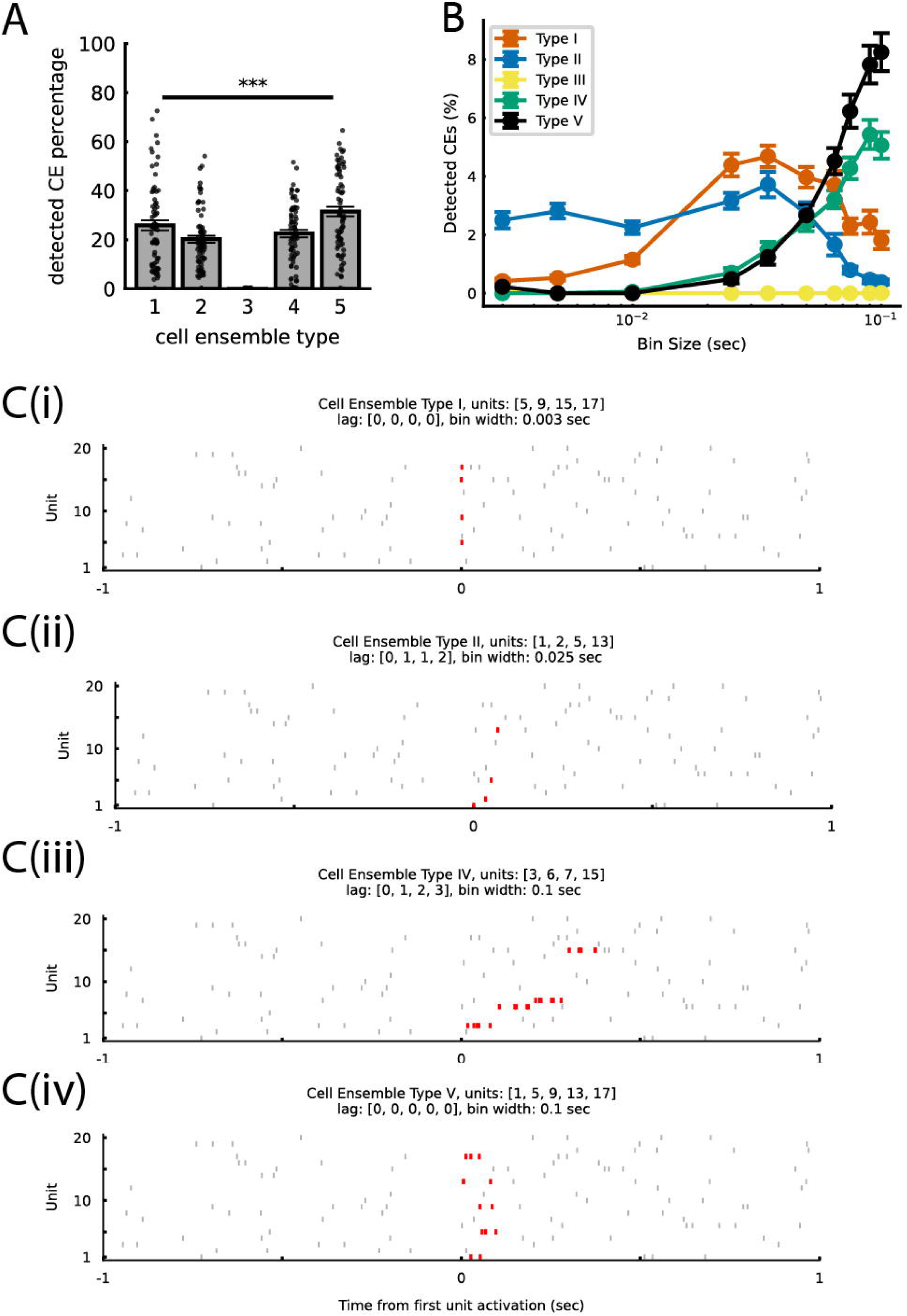
Cell ensemble types detected in rat primary motor cortex. **A.** Percentage of detected cell ensemble types according to Russo and Durstewitz (2017) (Kruskal–Wallis test, p = 9.83 × 10^-38^; mean ± sem; n = 60 sessions). **B.** Percentage of ensemble types across different bin sizes (mean ± sem; n = 60 sessions). **C.** Representative examples of detected cell ensembles. **C(i).** Type I (precise lag-0 synchronization with a single spike per neuron). **C(ii).** Type II (precise sequential pattern with a single spike per neuron). **C(iii)**. Type IV (rate pattern with temporal structure). **C(iv)**. Type V (simultaneous rate increase).

**Table 2.** Total number of detected cell ensembles during each recording session.

| Day | Rat 1 | Rat 2 | Rat 3 | Rat 4 |
| --- | --- | --- | --- | --- |
| 1 | 138 | 164 | 1904 | 132 |
| 2 | 362 | 172 | 3390 | 81 |
| 3 | 602 | 560 | 3426 | 635 |
| 4 | 217 | 549 | 9716 | 453 |
| 5 | 292 | 122 | 6976 | 504 |
| 6 | 1156 | 258 | 7741 | 65 |
| 7 | 873 | 225 | 2734 | 756 |
| 8 | 481 | 254 | 5298 | 313 |
| 9 | 1394 | 164 | 3557 | 95 |
| 10 | 2203 | 322 | 2995 | 156 |
| 11 | 2850 |  | 6095 | 199 |
| 12 | 1297 |  | 5121 | 114 |
| 13 | 1376 |  | 7035 | 136 |
| 14 | 588 |  | 8040 | 250 |
| 15 | 196 |  | 3908 |  |
| 16 |  |  | 4796 |  |
| 17 |  |  | 6905 |  |
| 18 |  |  | 3845 |  |
| 19 |  |  | 3696 |  |
| 20 |  |  | 3102 |  |

The types of detected cell ensembles varied depending on the bin size. Generally, cell ensembles with fine temporal structures, such as Types I, II, and III, tend to be detected in smaller bin sizes, whereas the ensembles with less clear temporal structures, such as Types IV and V, are more likely to be detected in larger bin sizes.

Consistent with this, Type II ensembles were detected from small to intermediate bin sizes (1-50 msec), and decreased for larger bin sizes (90-100 msec) (Figure 2B, blue). Types IV and V ensembles were not detected at small bin sizes (3-10 msec), but the detection increased from the intermediate (35 msec) to larger bin sizes, with the highest detection occurring at 100 msec (Figure 2B, green and black). Unlike our prediction that Type I ensembles should be detected at smaller bin sizes, they behaved a little differently, as their detection started at small bins (3-10 msec), peaked at intermediate bins (25-50 msec), and continued at larger bin sizes (65-100 msec) (Figure 2B, red). This is because Type I ensembles were identified as simultaneous activation of spikes within the same bin, which could happen at any bin size. However, since detection at larger bin sizes did not reflect precise spike synchronization, detections only in small bins should be considered as Type I ensembles. Representative examples of detected cell ensembles are shown in Figure 2C.

### Most cell ensembles were small and short-lived

The relative percentage of detected cell ensembles increased as the bin size increased (Figure 3A). The number of detected cell ensembles also grew as the number of recorded neurons in the experiment increased (Figure 3B), and the relationship was fitted as log (y) = 0.029x + 1.5459 (p = 1.80 x 10^-22^, r^2^ = 0.8084). The number of neurons in cell ensembles (population size) increased from 2-3 neurons for small bin sizes to 5-6 neurons for large bin sizes (Figures 3C(i) and 3D(ii). The matrices were normalized by the sum of all elements in Figure 3C(i) and by the column-wise sum in Figure 3C(ii). The results show that most cell ensembles were detected in larger bin sizes (Figure 3C(i)). We can also observe that cell ensembles in smaller bin sizes consist of fewer neurons, whereas those in larger bin sizes include a greater number of member neurons (Figure 3C(ii)). Analysis of the lifetime of cell ensembles revealed that most had brief activations, often lasting from zero lag to a few lags. (Figures 3D (i) and 3D (ii)). Figure 3D (i), where matrix elements were normalized by the sum of all elements, showed that 0- and short-lag cell ensembles are dominant at large bin sizes. Using normalization by column sum, Figure 3D (ii) indicated that cell ensembles with a lag of two are most common at small bin sizes, while ensembles tend to synchronize at larger bin sizes.

In summary, more cell ensembles were detected in larger bin sizes and in sessions with a higher number of recorded neurons. Most cell ensembles were small (2- 7 neurons) and short-lived (0-2 lags).

### Reach-related cell ensembles were grouped into four types based on their activation patterns

Among all detected cell ensembles, approximately 60% were activated during the reaching behavior. To characterize these reach-related cell ensembles, we analyzed their activation dynamics by K-means clustering. Clustering was performed separately on ten different bin sizes. For small bin sizes, the fluctuation of activation dynamics was large, and no clear clustering result was obtained. However, several stable patterns emerged in larger bin sizes, specifically for a bin size of 100 msec. Additionally, the largest percentage of cell ensembles was detected at 100 msec (Figure 3A). Since the bin size of 100 msec was also commonly used in memory reactivation studies (Wilson and McNaughton, 1994; Lee and Wilson, 2002; Carr et al., 2011), we focused on the clustering result at this bin size.

**Figure 3.**
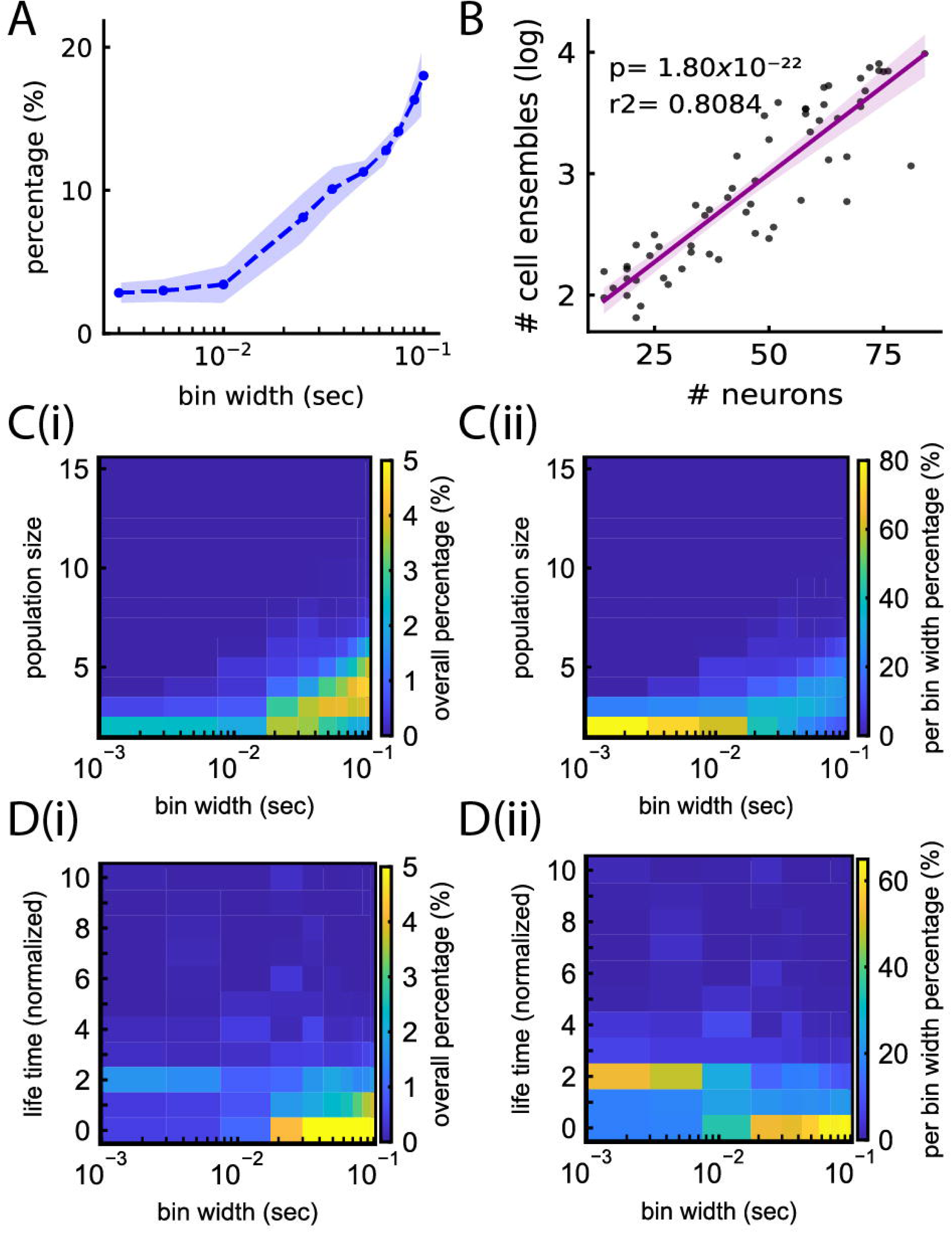
Properties of detected cell ensembles. **A.** Percentage of detected cell ensembles across different bin sizes (n=4 rats, mean ± sem). **B.** Relationship between the number of detected cell ensembles and the number of recorded neurons (n=60 sessions, scatter plots). Linear regression of the logarithm of ensemble number (y) against neuron number (x): log *y* = 0.021*x* + 1.5459, p = 1.80 x 10^-22^, r^2^=0.8084. **C.** Histogram of the ensemble’s population size (number of member neurons) for different bin sizes. **C(i).** The matrices were normalized by the sum of all elements. **C(ii).** The matrices were normalized by the column-wise sum. **D.** Histogram of the ensemble’s lifetime (lag between first and last neuron firing) for different bin sizes. **D(i).** The matrices were normalized by the sum of all elements. **D(ii).** The matrices were normalized by the column-wise sum.

Firstly, we evaluated the optimal number of clusters for each rat and each training day (Figure 4A). To test for any trends, the data were divided into three training phases: early (days 1-5), middle (days 6-10), and late (days 11-15). No significant difference was found across the training phases (Kruskal-Wallis test, p = 0.5397) and across animals (Kruskal-Wallis test, p = 0.62). To estimate the representative number of clusters, we created a histogram of the optimal K values and found that the mode was K = 4 (Figure 4B, Silhouette score at K = 4: 0.36 ± 0.01 across recording sessions). In the following analysis, K was fixed as 4.

**Figure 4.**
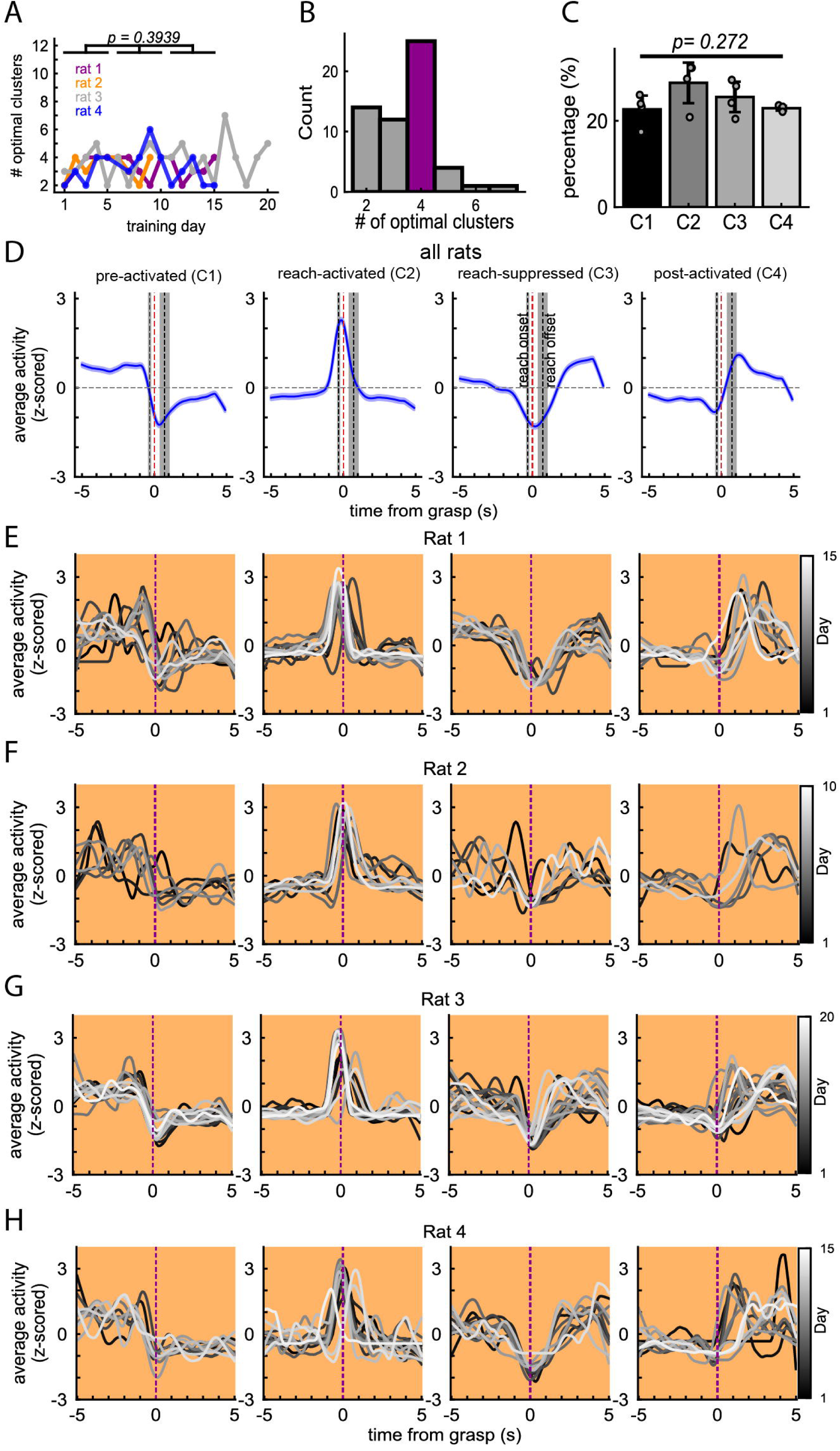
Clustering of reach-related cell ensembles. **A.** The optimal number of clusters (K) obtained by k-means clustering on individual sessions, evaluated with the Silhouette metric. Training days are separated into three groups, 1–5, 6–10, and 11–15 days. (Kruskal-Wallis test, across the training phases, p = 0.5397 and across animals, p = 0.62). **B.** Histogram of the optimal number of clusters. The mode (K = 4) is shown in magenta. **C.** Percentage of the cell ensembles assigned to each cluster (mean ± sd across n=4 rats, Kruskal-Wallis test, p= 0.272). **D.** Four clusters exhibiting distinct activation patterns around the reaching behavior. They are referred to as pre-activated (C1), reach- activated (C2), reach-suppressed (C3), and post-activated (C4). Blue lines and shaded areas represent the cell ensemble’s PETH and sem, respectively. The dotted red vertical line indicates grasping time. The gray band on the left shows the range of quantiles 1-3 for the reach-onset time relative to the grasping. The gray band on the right shows the range of quantiles 1-3 for the reach-offset time relative to the grasping time. **E-H.** Four reach-related clusters detected on different training days for each rat. Rat 1: Line color represents training days, from black (day 1) to white (day 15) **(E)**. Rat 2: Line color represents training days, from black (day 1) to white (day 10) **(F)**. Rat 3: Line color represents training days, from black (day 1) to white (day 20) **(G)**. Rat 4: Line color represents training days, from black (day 1) to white (day 15) **(H)**.

We found that the four different clusters exhibited distinct activation patterns during the reaching behavior (Figure 4D). The first cluster, C1, exhibited high activation before the reach onset. We call it ‘pre-activated’, and it may be related to preparatory activation. The second cluster, C2, exhibited a sharp increase in activation around the time of pellet grasping. We call it ‘reach-activated’, and it resembles the principal component activation pattern in the previous study (Eckert et al., 2020). The third cluster, C3, exhibited suppression around the time of pellet grasping, and we call it ‘reach-suppressed’. Finally, the fourth cluster, C4, exhibited high activation after the grasp, and we call it ‘post-activated’. No significant difference was found regarding the number of cell ensembles in each cluster type (Figure 4C, Kruskal-Wallis test, p=0.272). We also examined the activation patterns of the C1-C4 clusters for each animal, over training days (Figures 4E, 4F, 4G, and 4H correspond to rat 1, rat 2, rat 3, and rat 4, respectively. Relatively stable activation patterns were found throughout the training days.

In summary, the cell ensembles that were modulated around the grasping time were clustered into four types, each representing pre-activation (C1), reach-activation (C2), reach-suppression (C3), and post-activation (C4). Their activation patterns were similar among animals and remained stable throughout the training days.

### Reach-related cell ensembles replayed in both SWS and REM sleep

To investigate how the four types of reach-related cell ensembles were activated during rest, we compared their activation rates across all training days during the pre- task and post-task rest epochs. We found that the activation rates of all four types of reach-related cell ensembles were significantly higher during the post-task rest epoch, indicating that cell ensembles replayed after the task experience (Figure 5A (left, all rats): Wilcoxon signed-rank test: C1: n=47, p=6.66 ×10^-4^, C2: n=58, p=0.0055, C3: n=46, p=0.0073, C4: n=48, p=5.43 ×10^-5^). We repeated the analysis for the rapid and gradual learners separately but we didn’t observe a difference between them (Figure 5A (center, rapid learners): Wilcoxon signed-rank test, C1: n=40, statistic = 24, p= 10^-4^, C2: n=46, statistic= 12, p=1.6689 × 10^-8^, C3: n=36, statistic = 5, p=7.6293 × 10^-5^, C4: n=34, statistic =11, p=8 × 10^-4^. Figure 5A (right, gradual learners): Wilcoxon signed-rank test, C1: n=54, statistic = 23, p=9.5367 × 10^-6^, C2: n=70, statistic= 137, p=2.8328 × 10^-5^, C3: n=56, statistic = 51, p=2.4414 × 10^-5^, C4: n=62, statistic =42, p=1.045 × 10^-5^.

**Figure 5.**
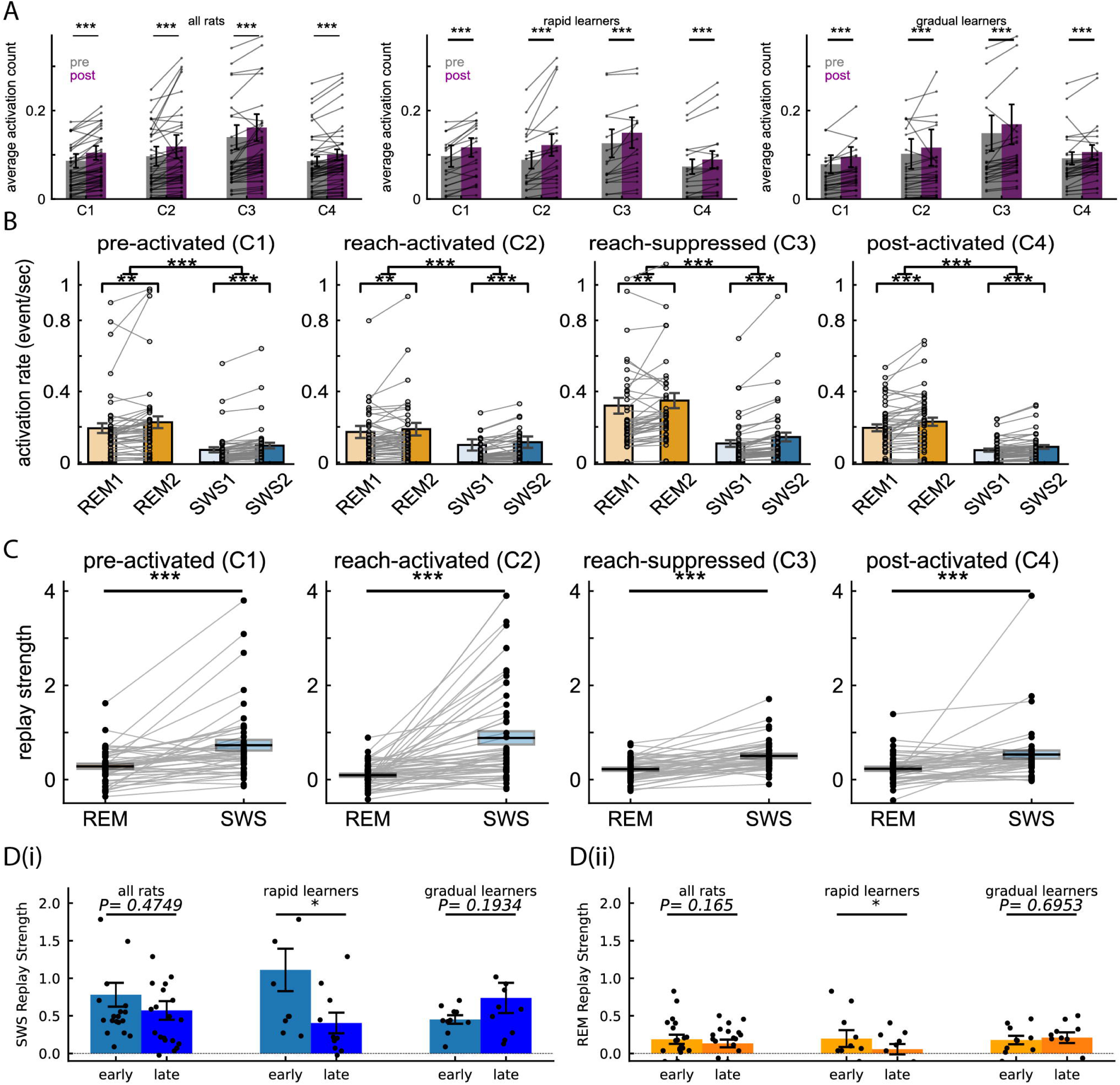
Replay of reach-related cell ensembles during SWS and REM sleep. **A.** Average activation counts of reach-related cell ensembles (C1, C2, C3, and C4) during pre-sleep and post-sleep. Activation counts were significantly higher during the post-task rest epoch (mean ± sem). **Left:** All rats combined. (Wilcoxon signed-rank test: C1: n=47, p=6.66 ×10^-4^, C2: n=58, p=0.0055, C3: n=46, p=0.0073, C4: n=48, p=5.43 ×10^-5^). **Middle:** Rapid learners. (Wilcoxon signed-rank test, C1: n=40, statistic = 24, p= 10^-4^, C2: n=46, statistic= 12, p=1.6689 × 10^-8^, C3: n=36, statistic = 5, p=7.6293 × 10^-5^, C4: n=34, statistic =11, p=8 × 10^-4^). **Right:** Gradual learners. (Wilcoxon signed-rank test, C1: n=54, statistic = 23, p=9.5367 × 10^-6^, C2: n=70, statistic= 137, p=2.8328 × 10^-5^, C3: n=56, statistic = 51, p=24414 × 10^-5^, C4: n=62, statistic =42, p=1.045 × 10^-5^). **B.** Activation rates (events/sec) during REM and SWS in the pre-task rest and post-task rest epochs (mean ± sem). **REM1-REM2 comparison**: REM2 activation rates were significantly higher than REM1 activation rates. (Wilcoxon signed-rank test: C1: n=47, p=6.66 ×10^-4^, C2: n=58, p=0.0055, C3: n=46, p=0.0073, C4: n=48, p=5.43 ×10^-5^). **SWS1-SWS2 comparison**: SWS2 activation rates were significantly higher than SWS1 activation rates. (Wilcoxon signed-rank test: C1: n=47, p=7.99 ×10^-8^, C2: n=58, p=6.10 ×10^-8^, C3: n=46, p=1.56 × 10^-12^, C4: n=48, p=1.22 ×10^-8^). **REM-SWS comparison**: REM activation rates were significantly higher than SWS activation rates. (Wilcoxon signed-rank test: C1: n=47, p=1.71 ×10^-8^, C2: n=58, p=9.33 ×10^-10^, C3: n=46, p=3.97 × 10^-13^, C4: n=48, p=1.47 ×10^-12^). **C.** Replay strength during REM and SWS. Replay was significantly stronger in SWS than REM (mean ± sem). (Wilcoxon signed-rank test; C1: n=47, p=7.97 ×10^-7^, C2: n=58, p=2.81 ×10^-9^, C3: n=46, p=2.70 × 10^-7^, C4: n=48, p=8.29 ×10^-6^). **D.** Replay strength during REM and SWS in the early (days 1–5) and the late (days 11–15; rat 2: days 6–10) training phases (mean ± sem). **D(i)**. SWS replay strength was analyzed for all rats combined (left), rapid learners (middle), and gradual learners (right). Wilcoxon signed-rank test: all rats, n = 40, p = 0.4304; rapid learners, n = 20, p = 0.0488; gradual learners, n = 20, p=0.1934. **D(ii)**, REM replay strength was analyzed for all rats combined (left), rapid learners (middle), and gradual learners (right): Wilcoxon signed-rank test: all rats, n = 40, p=0.165; rapid learners, n = 20, p=0.0371; gradual learners, n = 20, p=0.6953.

Next, we assessed the activation rates for SWS and REM sleep (SWS1 and REM1 in the pre-task rest epoch and SWS2 and REM2 in the post-task rest epoch). The activation rates were significantly higher during the post-task rest epoch compared to the pre-task rest epochs for both REM and SWS, and the overall activation rates were significantly higher in REM than in SWS (Figure 5B, Table 3). However, when the replay was evaluated using the replay strength, replay was significantly stronger in SWS than REM (Figure 5C, Wilcoxon signed-rank test, C1: n=47, statistic = 116, p=5.58 × 10^-7^, C2: n=58, statistic= 98, p=4.49 × 10-9, C3: n=46, statistic = 105, p=2.11 × 10^-7^, C4: n=48, statistic = 177, p=8.29 x 10^-6^). These results suggest that while the total replay of reach-related cell ensembles was greater in REM than in SWS, the normalized increase in replay was higher in SWS than in REM.

**Table 3.** Statistical results of activation rate comparison across different sleep stages (Wilcoxon signed-rank test, Figure 5A).

| Reach group | REM1 vs REM2 |  | SWS1 vs SWS2 |  | REM vs SWS |  |
| --- | --- | --- | --- | --- | --- | --- |
|  | statistic | pvalue | statistic | pvalue | statistic | pvalue |
| C1 (n=47) | 216 | $6.66 \times 10^{-4}$ | 42 | $7.99 \times 10^{-8}$ | 18 | $1.71 \times 10^{-8}$ |
| C2 (n=58) | 440 | 0.0055 | 156 | $6.10 \times 10^{-8}$ | 65 | $9.33 \times 10^{-10}$ |
| C3 (n=46) | 298 | 0.0073 | 11 | $1.56 \times 10^{-12}$ | 6 | $3.97 \times 10^{-13}$ |
| C4 (n=48) | 211 | $5.43 \times 10^{-5}$ | 88 | $1.22 \times 10^{-8}$ | 17 | $1.47 \times 10^{-12}$ |

We also examined whether the replay depended on the training phases. Firstly, we combined all four rats and compared the average replay strength between the early training phase (Days 1-5 for all animals) and the late training phase (Days 10-15 for rats 1, 3, and 4; Days 5-10 for rat 2). The average replay strength was higher in the early phase than in the late phase for both SWS and REM, but neither of the differences was statistically significant (Figures 5D(i) and 5D(ii). For SWS, the Wilcoxon signed-rank test, p=0.4749; For REM, the Wilcoxon signed-rank test, p=0.165).

Next, we divided the data into the fast and gradual learners and repeated the analysis. The rapid learners exhibited significantly more replay in the early training phase than in the late training phase, in both SWS and REM sleep (Figures 5D(i) and 5D(ii). For SWS, Wilcoxon signed-rank test, p=0.0488; For REM, Wilcoxon signed-rank test, p=0.0371). On the other hand, the gradual learners showed the opposite pattern but the difference was not statistically significant (Figures 5D(i) and 5D(ii)). For SWS, Wilcoxon signed-rank test, p=0.1934; For REM, Wilcoxon signed-rank test, p=0.6953).

In summary, replay of reach-related cell ensembles was detected in both SWS and REM sleep. The rapid learners exhibited significantly more replay during the early training phase, which coincided with their rapid performance gains. The gradual learners exhibited a non-significant increase in replay in the late training phase.

### Reach-related cell ensembles replayed preferentially during SO-spindles

Previous investigations using PCA have demonstrated that memory reactivation occurs during spindles in SWS (Ramanathan et al., 2015; Eckert et al., 2020). Here, we also examined whether spatio-temporal patterns were preferentially replayed during spindles. We compared the average activation count between spindle and non-spindle periods in post-SWS and found that all four types of reach-related cell ensembles became significantly more active during the spindles than between them. (Figure 6A(i), Wilcoxon signed-rank test, C1: n=47, statistic=141, p=1.61 × 10^-6^, C2: n=58, statistic=50, p=1.05 × 10^-9^, C3: n=46, statistic=182, p=4.04 ×10^-5^, C4: n=48, statistic=155, p=1.53 × 10^-5^). We also tested for a difference between rapid and gradual learners. Both groups showed significantly stronger activation during spindles than non-spindles, except for the reach-suppressed (C3) and post-activated (C4) ensembles in rapid learners. (Figure 6A(ii), rapid learners, Wilcoxon signed-rank test, C1: statistic=39, p=0.012, C2: statistic=9, p=7.86 × 10^-6^, C3: statistic=50, p=0.1297, C4: statistic=27, p=0.0637; Figure 6A(iii), gradual learners, Wilcoxon signed-rank test, C1: statistic= 21, p= 10^-4^, C2: statistic=52, p=7.05 × 10^-5^, C3: statistic=2, p= 8.94 × 10^-8^, C4: statistic=27, p=3.75 × 10^-5^).

**Figure 6.**
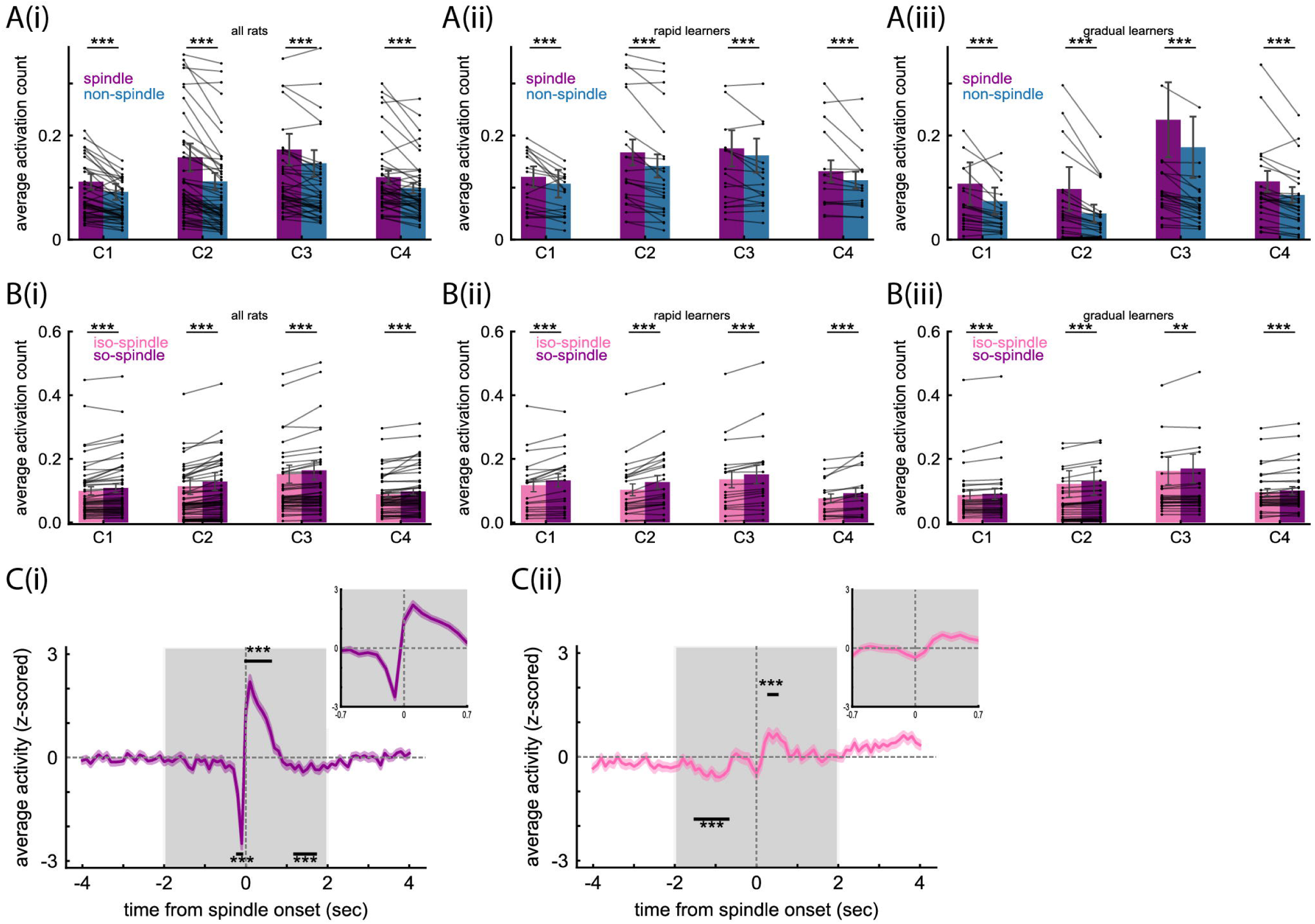
Replay of reach-related cell ensembles during sleep spindles. **A.** Average activation counts of reach-related cell ensembles (C1, C2, C3, and C4) during spindles and non-spindles (mean ± sem). **A(i)**. All rats combined. Activation counts were significantly higher during spindles than non-spindles (Wilcoxon signed-rank test, C1: n=47, statistic=141, p=1.61 × 10^-6^, C2: n=58, statistic=50, p=1.05 × 10^-9^, C3: n=46, statistic=182, p=4.04 ×10^-5^, C4: n=48, statistic=155, p=1.53 × 10^-5^). **A(ii)**. Rapid learners. Activation counts of the pre-activated (C1) and reach-activated (C2) ensembles were significantly higher during spindles than during non-spindles. The reach-suppressed (C3) and post-activated (C4) did not show a significant difference (Wilcoxon signed-rank test, C1: statistic=39, p=0.012, C2: statistic=9, p=7.86 × 10^-6^, C3: statistic=50, p=0.1297, C4: statistic=27, p=0.0637). **A(iii)**. Gradual learners. Activation counts were significantly higher during spindles than non-spindles (Wilcoxon signed-rank test, C1: statistic= 21, p= 10^-4^, C2: statistic=52, p=7.05 × 10^-5^, C3: statistic=2, p= 8.94 × 10^-8^, C4: statistic=27, p=3.75 × 10^-5^). **B.** Average activation counts of reach-related cell ensembles (C1, C2, C3, and C4) in isolated spindles and SO-spindles during the post-task epoch (mean ± sem). **B(i)**. All rats combined. Activation counts were significantly higher during SO-spindles than isolated spindles (Wilcoxon signed-rank test, C1: statistic=95, p= 1.12 × 10^-6^, C2: statistic =84.5, p= 5.86 × 10^-8^, C3: statistic=105, p= 8.78 ×10^-6^, C4: statistic= 134.5, p= 5.45 × 10^-6^). **B(ii)**. Rapid learners. Activation counts were significantly higher during SO-spindles than isolated spindles (Wilcoxon signed-rank test, C1: statistic=24, p= 8.85× 10^-4^, C2: statistic =12, p= 1.66×10^-5^, C3: statistic=5, p= 7.62 ×10^-5^, C4: statistic= 11, p= 8 × 10^-4^). **B(iii)**. Gradual learners. Activation counts were significantly higher during SO-spindles than isolated spindles (Wilcoxon signed-rank test, C1: statistic=23, p= 9.53 × 10^-6^, C2: statistic =137, p= 0.0028, C3: statistic=51, p= 2 ×10^-4^, C4: statistic= 42, p= 1.04 × 10^-5^). **C.** Peri-event time histograms (PETHs) of the activation of reach-related cell ensembles around spindle onset in post-task epoch (n=60, mean ± sem). **C(i)**. PETH around the SO-spindle onset **C(ii)**. PETH around the isolated spindle onset. In both panels, the horizontal black line indicates periods where activation was significantly different from baseline, which was defined as the mean activation during (-4, -2) sec and (2, 4) sec relative to spindle onset (Wilcoxon signed-rank test, Bonferroni-corrected; *** p < 0.001).

Next, we examined whether reach-related cell ensembles were more strongly activated during SO-spindles (spindles that followed slow oscillations) than isolated spindles. We confirmed that this is the case for all rats (Figure 6B(i), Wilcoxon signed-rank test, C1: statistic=95, p= 1.12 × 10^-6^, C2: statistic =84.5, p= 5.86 × 10^-8^, C3: statistic=105, p= 8.78 ×10^-6^, C4: statistic= 134.5, p= 5.45 × 10^-6^), but we did not observe a difference between the rapid and gradual learners (Figure 6B(ii), Rapid learners, Wilcoxon signed-rank test, C1: statistic=24, p= 8.85× 10^-4^, C2: statistic =12, p= 1.66×10^-5^, C3: statistic=5, p= 7.62 ×10^-5^, C4: statistic= 11, p= 8 × 10^-4^; Figure 6B(iii), gradual learners, Wilcoxon signed-rank test, C1: statistic=23, p= 9.53 × 10^-6^, C2: statistic =137, p= 0.0028, C3: statistic=51, p= 2 ×10^-4^, C4: statistic= 42, p= 1.04 × 10^-5^).

To gain insights into the temporal evolution of reach-related cell ensemble activations, we calculated PETHs around the onset of spindles. For the SO-spindles, the activation was significantly suppressed just before the spindle onset but increased sharply after the onset. (Figure 6C(i). Wilcoxon signed-rank test, p<0.001, Bonferroni corrected). This is likely to correspond to the down-to-up state transition during slow oscillations (Steriade et al., 1993; Cash et al., 2009). For the isolated spindles, a much weaker suppression and increase of activations were observed compared to the SO-coupled spindles (Figure 6C(ii). Wilcoxon signed-rank test, p<0.001, Bonferroni corrected). Thus, we confirmed that reach-related cell ensembles were more strongly activated during SO-spindles than isolated spindles.

### Reach-related cell ensembles were modulated during SWRs

We investigated the relationship between hippocampal SWRs and the activation of the reach-related cell ensembles by comparing the activation during SWRs with that during randomly selected periods between SWRs. About 30% of reach-related cell ensembles were modulated during SWRs and the rapid and gradual learners showed opposite modulation trends. The cell ensembles exhibited three distinct activation patterns around SWRs: enhanced modulation, suppressive modulation, and non-significant modulation (Figure 7A). The enhanced cell ensembles showed increased activity during SWRs, peaking just before the center of the SWRs. The suppressed cell ensembles exhibited a plateau of inhibition during SWRs. The proportions of the enhanced, suppressive, and non-significantly modulated cell ensembles were 12.64% ± 1.30%, 16% ± 1.47%, and 71.34% ± 1.30%, respectively (Figure 7B). Significant differences were observed between the enhanced and non-significant modulations, as well as between the suppressive and non-significant modulations. No significant difference was found between the enhanced and suppressive modulations (Friedman test, statistics= 87.483, p=1 × 10^-19^, Pairwise Wilcoxon signed-rank tests post hoc test, Bonferroni correction).

**Figure 7.**
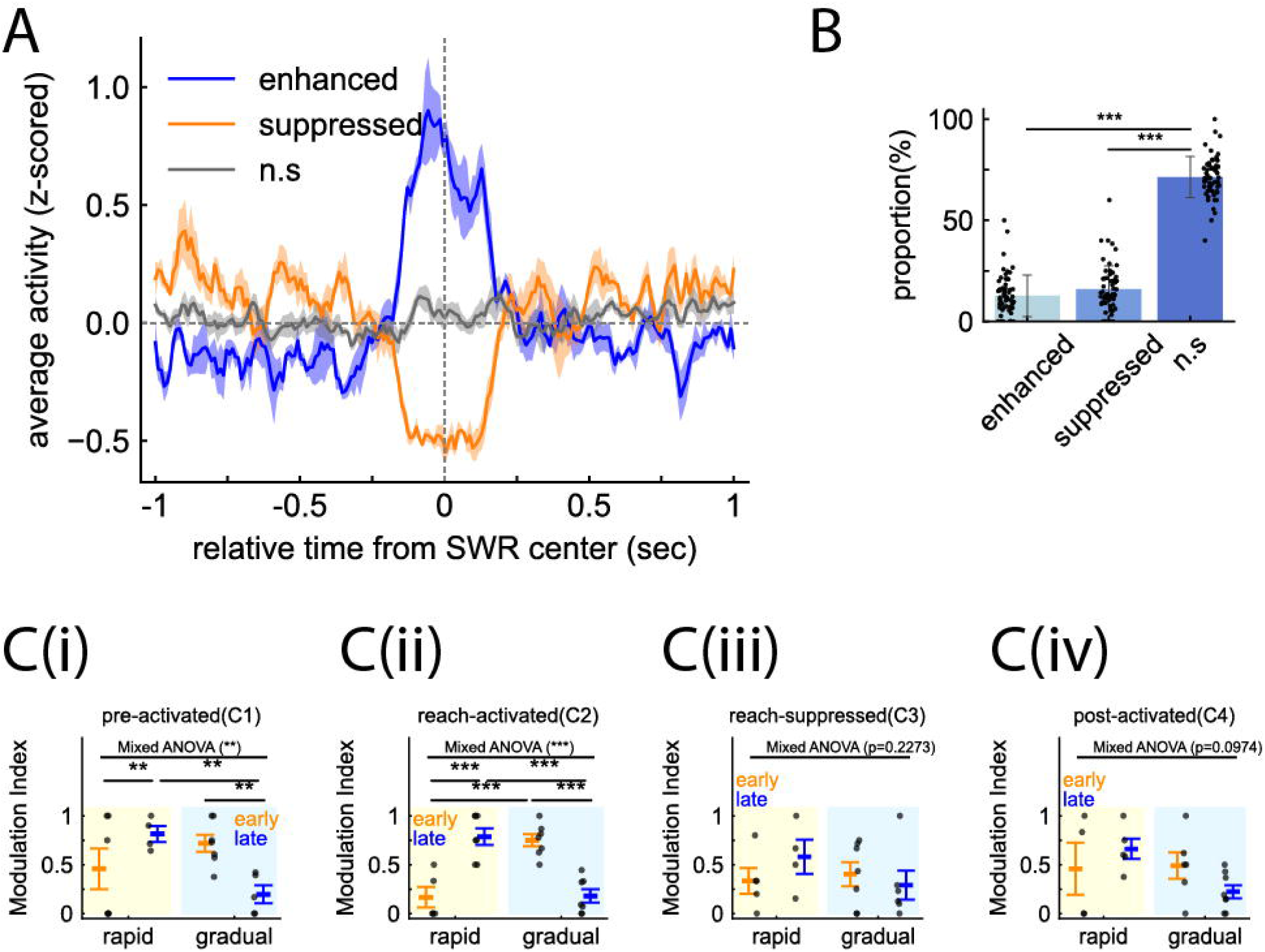
Modulation of reach-related cell ensembles during SWRs. **A.** Reach-related ensembles exhibited distinct temporal patterns around SWRs. Blue, orange, and gray lines are enhanced modulation, suppressed modulation, and non-significant modulation (ns), respectively (mean ± sem). **B.** Percentage of the SWR-modulated reach-related cell ensembles. The proportions of the enhanced, suppressive, and non-significantly modulated cell ensembles are 12.64% ± 1.30%, 16% ± 1.47%, and 71.34% ± 1.30%, respectively (mean ± SD). (Kruskal-Wallis test, n=60, p=7.84 × 10^-27^, Dunn’s post hoc test with Bonferroni correction). **C.** SWR modulation of the four reach-related cell ensembles (C1-C4) during early and late training phases (mean ± sem). Two-way mixed design ANOVA, Interaction effect, significant post-hoc t-test (Bonferroni corrected), ** 0.001<p<0.01, ***p<0.001. **C(i)**. Pre-activated cell ensembles (C1) (F(1,18)=9.8226, p=0.0057). **C(ii)**. Reach-activated cell ensembles (C2) (F(1,24)= 53.85, p=1.82 ×10^-7^). **C(iii)**. Reach-suppressed cell ensembles (C3) (F(1,19)=1.5565, p=0.2273). C(iv). Post-activated cell ensembles (C4) (F(1,19)=3.0393, p=0.0974).

We also examined the relationship between the animals’ learning types (fast and gradual learners) and the modulation of activation during SWRs. We calculated the modulation index, defined as the ratio of SWR-enhanced ensembles to the total of SWS- enhanced and SWS-suppressed ensembles (see methods). The rapid learners exhibited a tendency for cell ensembles to be suppressed (modulation index < 0.5) in the early training phase, whereas they were enhanced (modulation index > 0.5) in the late training phase. In contrast, the gradual learners showed the opposite pattern, with cell ensembles being enhanced in the early training phase and suppressed in the late training phase (Figure 7C). Then, a 2×2 mixed-design ANOVA was conducted with the training phase as a within-subject factor and the learning type as a between-subject factor, followed by a post-hoc t-test (Bonferroni corrected). We found a significant interaction between the rapid and gradual learners for the pre-activated and reach- activated ensembles, but not for the reach-suppressed and post-activated ensembles (Pre-activated (C1): Rat type effect: F=0.5025, p=0.4874, Phase effect: F=0.816, p=0.3782, Interaction effect: F=9.8226, p=0.0057; Reach-activated (C2): Rat type effect: F=0.6779, p=0.4191; Phase effect: F=0.14, p=0.7093; Interaction effect: F=53.85, p=1.82x10^-7^; Reach-suppressed (C3): Rat type effect: F=0.4072, p=0.5313, Phase effect: F=0.0579, p=0.8124, Interaction effect: F=1.5565, p=0.2273; Post-activated: Rat type effect: F=3.0267, p=0.098, Phase effect: F=0.3882, p=0.5406, Interaction effect: F=3.0393, p=0.0974).

In summary, about 30% of reach-related cell ensembles were modulated during SWRs, indicating the involvement of the hippocampus. The rapid and gradual learners exhibited opposite modulation trends, with significant differences found for the pre-activated and the reach-activated ensembles.

### SWRs and SO-spindles co-occurred and the overall coupling strength changed in opposite directions for rapid and gradual learners

As was explored for explicit memory, the temporal coupling of SWRs and spindles is important for communication between the hippocampus and the cortex (Siapas and Wilson, 1998; Maingret et al., 2016; Pedrosa et al., 2024). Therefore, we examined the timing relationship between the SO-spindle onset and the SWR center during the post-task rest epoch (rest 2). We focused on the SO-spindle onset because more replay occurred in the early phase of SO-spindles (Figure 6C). We generated the PETHs within ±2 seconds of the SO-spindle onset, separately for the early and late phases of training. We showed the results with four rats (Figure 8A(i)), rapid learners only (Figure 8A(ii)), and gradual learners only (Figure 8A(iii)).

**Figure 8.**
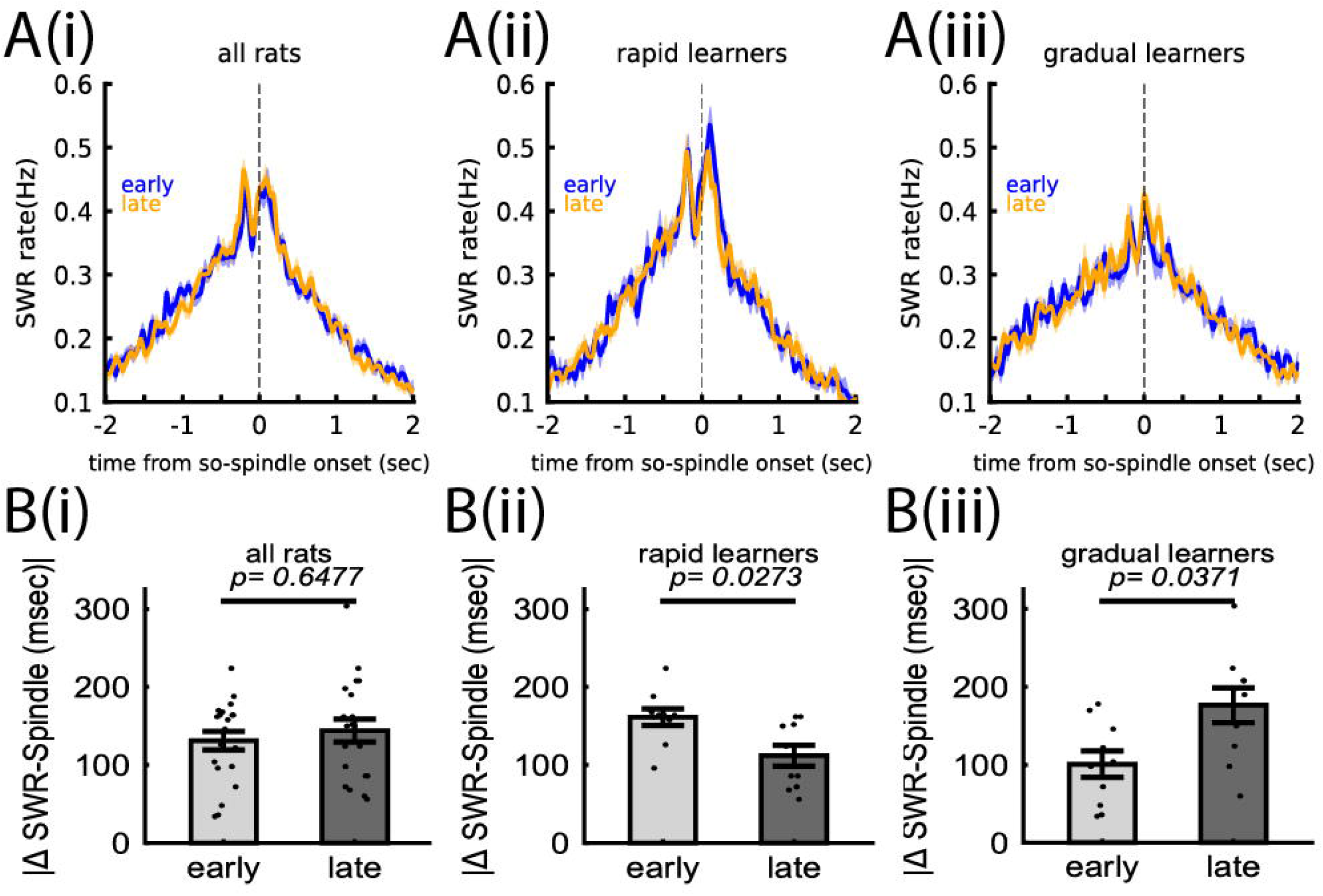
Temporal relationship between SWRs and SO-spindles. **A.** PETH of SWRs that is aligned to SO–spindle onset. PETH was calculated for early and late training phases separately (mean ± sem, blue: early phase, orange: late phase). **A(i)**. All rats. **A(ii)**. Rapid learner. **A(iii)**. Gradual learners. **B.** Absolute value of temporal lag between SWR peaks and SO–spindle onset, calculated for early and late training phases separately (mean ± sem). **B(i)**. All rats (Wilcoxon signed-rank test, n=40 sessions, statistic= 92.5, p=0.6477). **B(ii)**. Rapid learners (Wilcoxon signed-rank test, n=10 sessions, n=6, p=0.0273). **B(iii)**. Gradual learners (Wilcoxon signed-rank test, n=10 sessions, n=7, p=0.0371).

A triangular shape with a peak around lag-zero indicated a tendency for SWRs and SO-spindles to co-occur. The strength of temporal coupling was examined by calculating the separation of the dual central peaks from lag-zero. To estimate the overall tendency, we used the absolute value, without distinguishing whether the peaks occurred before or after the lag-zero. With four rats included in the analysis, no statistical difference was observed between the early and late phases of training (Figure 8B(i), Wilcoxon signed-rank test, p = 0.6477). However, the temporal separation for the rapid learners significantly decreased from the early to late phase of training (Figure 8B(ii), Wilcoxon signed-rank test, p = 0.0273). In contrast, the temporal separation for the gradual learners significantly increased from the early to late phase of training (Figure 8B(iii), Wilcoxon signed-rank test, p=0.0371).

In summary, SWRs and SO-spindles tended to co-occur during the rat’s skilled reaching learning. Over the course of training, the overall strength of coupling changed in opposite directions for the rapid and gradual learners.

## Discussion

We analyzed spike data from the rat’s primary motor cortex while the animals learned the skilled reaching task and rested. A combination of unsupervised cell ensemble detection and clustering revealed that the reaching-related cell ensembles could be categorized into four types, and that they replayed during SWS and REM sleep, specifically during SO-spindles. About 30% of them were modulated during the hippocampal SWRs, and the overall direction of modulation depended on the phase of training (early or late) and whether the animals were fast or gradual learners.

### Size of detected cell ensembles

Approximately 40% of detected cell ensembles showed synchronous or precise sequential activations (Types I and II), while about 60% exhibited firing rate modulations (Types IV and V) (Figure 2A). The size of the detected cell ensembles was relatively small, typically 2-3 neurons for the smaller bins and 4-6 neurons for the larger bins (Figure 3D). The sequential activation of small cell ensembles may provide a building block for precise sequence computation, supporting chunking-based information processing (Graybiel, 1998; Peters et al., 2014). They also align with the idea of precise coordination of neural activities, such as synfire chains (Abeles et al., 1993; Ikegaya et al., 2004). At the same time, it should be noted that the employed unsupervised method, which applied strict statistical significance tests at each agglomerative step, could result in detecting smaller cell ensembles. Interestingly, our results are comparable to those obtained using PCA or ICA, where only a small subset of neurons shows significantly high contributions to the principal or independent components (van de Ven et al., 2016; Kim et al., 2019). In summary, it is important to recognize that different detection methods that employ different computational approaches could produce different detection results. Thus, further analysis using different computational criteria will add more insight into the activation of cell ensembles.

### Stability and replay of reach-related cell ensembles

This study revealed the activation of four types of cell ensembles related to reaching behavior (C1-C4 ensembles). This richer neural activation pattern aligns with earlier research reporting multiple neural activities linked to motor learning tasks (Rokni et al., 2007; Peters et al., 2014; Peters et al., 2017). Their highly consistent activation across training days was somewhat surprising (Figures 4B-4F), but it may indicate the presence of fundamental neural motifs underlying rats’ skilled reaching movements (Weinreb et al., 2024). We also detected replay of all four reach-related cell ensembles, including the reach-suppressed cell ensembles (C3). It may be related to inhibitory neural activities that are crucial for stabilizing motor circuits by helping stabilize the networks through memory replay (Eisenstein et al., 2023). In contrast to REM preactivation reported in our previous study (Eckert et al., 2020), this study observed REM replay (Figure 5). One possible explanation is the difference in analysis methods: Eckert’s study used PCA, where principal components reflect overall coincident neural activities but are not sensitive to spatio-temporal patterns. Instead, the unsupervised method used in this study can detect a broader range of cell ensemble activities. It is also worth noting that the brain may utilize different neural representations simultaneously (Riehle et al., 1997). Examining brain signals with multiple analysis methods could reveal different aspects of neural information processing.

### SO-spindle coupling and the interaction between the motor cortex and the hippocampus

Replay during SWS preferentially occurred during SO-spindles (Figure 6). Biologically, SOs are associated with the down-to-up transition and can reset cortical excitability (Massimini et al., 2004), whereas spindles are associated with synaptic plasticity (Rosanova and Ulrich, 2005). The temporal coupling of spindles with SOs is likely to induce long-term potentiation in the motor cortical circuits, facilitating the integration and stabilization of new memories (Darevsky et al., 2024). Thus, the interplay between SOs and spindles, which aligns bursts of cell ensemble activation during the up state, can be beneficial for memory consolidation during sleep.

Unlike explicit memory, the extent to which learning of implicit memory depends on the hippocampus has been a subject of debate (Squire, 2004; Degonda et al., 2005; Steinkrauss and Slotnick, 2024). Our results support the involvement of the hippocampus in rats’ skill learning, where about 30% of reach-related cell ensembles were either enhanced or suppressed during SWRs (Figure 7). Interestingly, the rapid learners exhibited enhanced modulation and tighter coupling of SWRs and SO-spindles in the late training phase, whereas these were observed in the early training phase for the gradual learners (Figures 7 and 8). More communication between the primary motor cortex and the hippocampus may occur at different training phases for the rapid and gradual learners.

### Hippocampal involvement in rats’ skill learning

Strong replay for the rapid learners occurred when they exhibited significant performance improvement during the early training phase, whereas strong replay for the gradual learners occurred when they continued to improve performance in the late training phase (Figures 1 and 5). Interestingly, their initial performance differed significantly, with approximately 15% and 45% success rates for the rapid and gradual learners, respectively (Figure 1). Additionally, the amount of replay of the rapid learners during the late training phase was similar to that of the gradual learners during the early training phase (Figure 5). Taken together, these findings suggest that the late training phase of rapid learners and the early training phase of gradual learners may be similar. Based on these observations and the discussions above, we propose a hypothetical diagram of rats’ skill learning, including the changes in success rate, replay strength, and the hippocampus involvement, for both fast and gradual learners (Figure 9).

**Figure 9.**
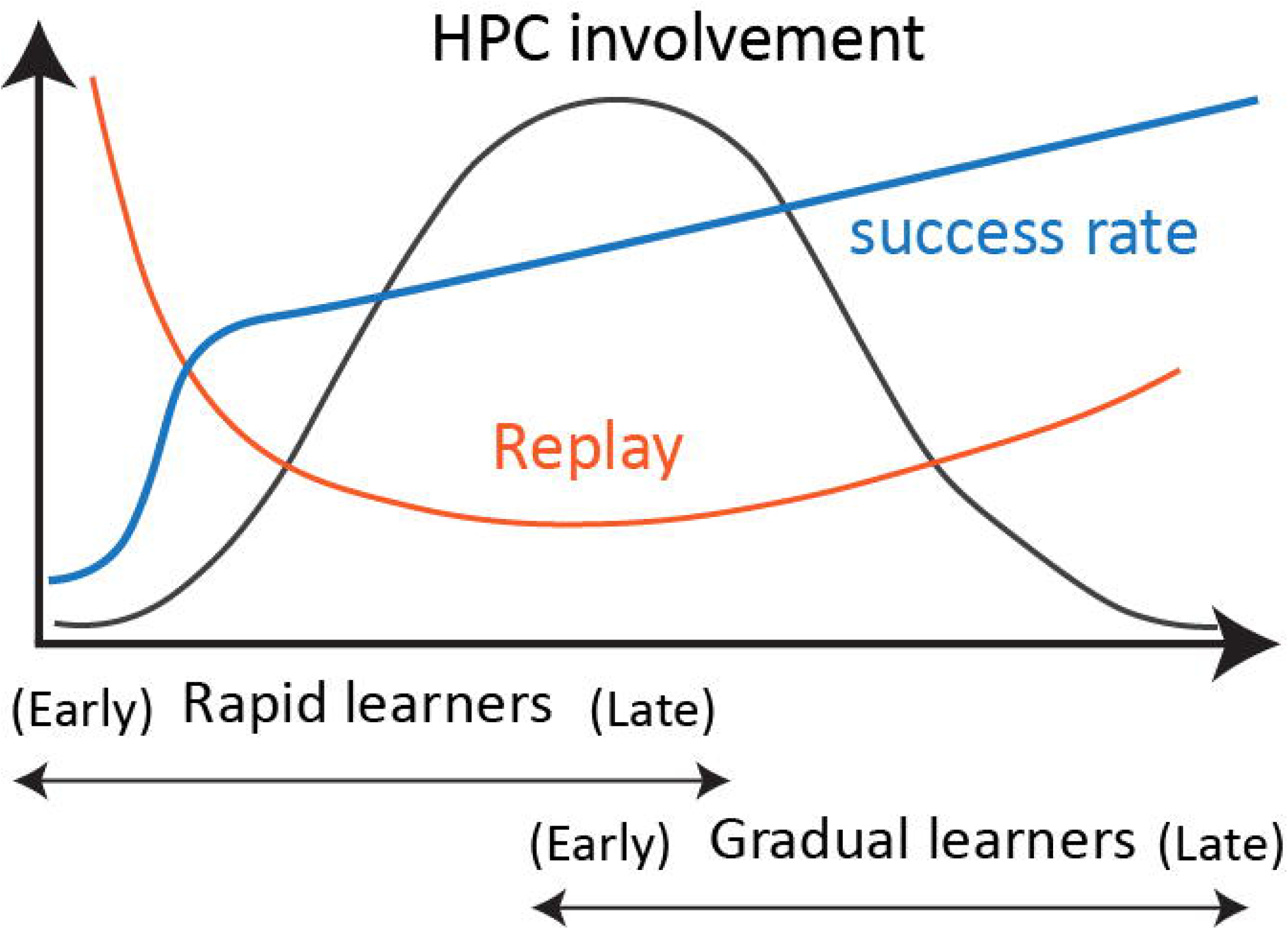
Hypothetical model illustrating possible cortico-hippocampal communications during sleep for rats’ motor skill learning. The blue curve represents the improvement in the success rate. The red curve represents the change in replay strength. The black curve represents the involvement of the hippocampus in skill learning. All three curves are smoothly connected during the late training phase of rapid learners and the early training phase of gradual learners.

The improvement in the success rate is represented by the blue curve. The rapid learners begin with a low success rate, but exhibit rapid improvement during the early training phase, followed by slow improvement during the late training phase.

Considering the similar success rate during the early training phase of the rapid learners and the late training phase of the gradual learners (Figure 1), the blue curve is smoothly connected in the middle of the figure.

The change in replay strength is represented by the red curve. The rapid learners exhibit strong replay during the early training phase, and it decreases to a moderate level in the late phase. Based on the similarity of replay strength during the early training phase of the rapid learners and the late training phase of the gradual learners (Figure 5), the red curve is smoothly connected in the middle of the figure.

The hypothetical involvement of the hippocampus is represented by the black curve. During the early training phase, cortical ensembles in the rapid learners are suppressed during SWRs (Figure 7) and loosely coupled to SWRs and SO-spindles (Figure 8), suggesting diminished hippocampal influence. During the late phase, these relationships are reversed. Ensemble activity is enhanced during SWRs (Figure 7) and the coupling of SWRs and SO-spindles is tightened (Figure 8), suggesting that the influence of the hippocampus is increased. Regarding the gradual learners, ensembles exhibit the opposite pattern. From the early to late training phases, replay modulation shifts from enhanced to suppressed, and the timing relationship of SWRs and SO-spindles moves from tight to loose coupling. The black curve is smoothly connected at the center of the figure, forming a bell-shaped curve for the hippocampal involvement.

In summary, rapid learners may initially focus on skill improvement without much hippocampal influence. Then, after gaining basic skill proficiency, they might refine reaching skills using hippocampal activity that carries spatial/contextual information. Conversely, the gradual learners, who demonstrate decent reaching performance from the start, might initially rely on hippocampal information. Then, hippocampal involvement seems to diminish as they continue to learn reaching skills.

### Limitations

This study has several limitations. First, due to the nature of unsupervised detection, the relationship between cell ensemble activations and behavior remains elusive. Second, because of the technical difficulties, we have not examined how cell ensembles detected at one bin size (for example, 100 msec during a task) become active at a different bin size (for example, 10 msec during rest). However, this is crucial because replay can be temporarily compressed. Third, SOs, spindles, and SWRs can also occur during quiet wakefulness. Future studies should investigate the relationship between SOs and spindles and how the hippocampal and cortical regions interact during quiet wakefulness. Fourth, because of mathematical challenges, the ensemble detection method used cannot track the changes in cell ensemble configurations over time.

However, since neuroplasticity and changes in cell ensembles are hallmarks of the brain, future research should address this problem.

## Contributions

M. Tatsuno conceived and designed the project. M. Eckert conducted the experiment and collected data. P. Nazari performed the analysis. M. Tatsuno and M. Eckert helped with the discussion of analysis and results. P. Nazari and M. Tatsuno wrote the paper with help from M. Eckert.

## Conflict of interest statement

The authors declare no competing financial interests.

## Acknowledgments

This work was supported by the Natural Sciences and Engineering Research Council of Canada (NSERC) 2020-06342 awarded to M.T., and was partly enabled by support from the Digital Research Alliance of Canada. We thank Bruce L. McNaughton and Javad Karimi Abadchi for helpful comments.

## Notes

### Competing Interest Statement

The authors have declared no competing interest.

