## supplemental tables for "Behaviorally relevant cell ensembles in rat motor cortex are replayed during sleep and implicate hippocampal involvement in motor skill learning"

**Supplemental Materials**

Supplemental Table 1. Number of detected cell ensembles per rat.

Rat 1

| Bin Size (msec) | Day1 | Day2 | Day3 | Day4 | Day5 | Day6 | Day7 | Day8 | Day9 | Day10 | Day11 | Day12 | Day13 | Day14 | Day15 |
|---|---|---|---|---|---|---|---|---|---|---|---|---|---|---|---|
| 3 | 7 | 18 | 10 | 9 | 23 | 49 | 31 | 22 | 51 | 39 | 68 | 45 | 46 | 19 | 12 |
| 5 | 6 | 15 | 14 | 12 | 18 | 54 | 19 | 12 | 39 | 41 | 128 | 81 | 53 | 26 | 8 |
| 10 | 9 | 15 | 21 | 14 | 27 | 38 | 22 | 15 | 36 | 69 | 277 | 107 | 88 | 14 | 10 |
| 25 | 14 | 30 | 60 | 37 | 41 | 63 | 51 | 36 | 90 | 186 | 380 | 195 | 147 | 56 | 15 |
| 35 | 14 | 38 | 58 | 34 | 46 | 88 | 94 | 55 | 114 | 194 | 455 | 230 | 191 | 75 | 19 |
| 50 | 14 | 39 | 74 | 20 | 32 | 111 | 105 | 63 | 134 | 219 | 252 | 113 | 141 | 64 | 29 |
| 65 | 15 | 43 | 87 | 21 | 27 | 143 | 118 | 68 | 172 | 248 | 243 | 103 | 141 | 70 | 23 |
| 75 | 16 | 51 | 84 | 23 | 24 | 202 | 128 | 68 | 200 | 323 | 329 | 135 | 172 | 81 | 27 |
| 90 | 20 | 58 | 100 | 25 | 28 | 192 | 140 | 71 | 281 | 423 | 367 | 134 | 185 | - | 25 |
| 100 | 23 | 55 | 94 | 22 | 26 | 216 | 165 | 71 | 277 | 461 | 351 | 154 | 212 | 183 | 28 |

Rat 2

| Bin Size (msec) | Day1 | Day2 | Day3 | Day4 | Day5 | Day6 | Day7 | Day8 | Day9 | Day10 |
|---|---|---|---|---|---|---|---|---|---|---|
| 3 | 6 | 12 | 16 | 15 | 16 | 11 | 9 | 2 | 4 | 16 |
| 5 | 7 | 10 | 15 | 12 | 15 | 6 | 8 | 8 | 5 | 14 |
| 10 | 7 | 5 | 11 | 13 | 4 | 6 | 6 | 5 | 5 | 10 |
| 25 | 10 | 13 | 44 | 37 | 10 | 13 | 19 | 15 | 6 | 20 |
| 35 | 12 | 13 | 50 | 46 | 13 | 24 | 21 | 24 | 17 | 26 |
| 50 | 19 | 18 | 58 | 66 | 15 | 25 | 30 | 26 | 18 | 38 |
| 65 | 23 | 24 | 75 | 77 | 11 | 31 | 34 | 45 | 20 | 40 |
| 75 | 21 | 19 | 87 | 78 | 13 | 37 | 28 | 40 | 24 | 49 |
| 90 | 32 | 28 | 100 | 100 | 12 | 50 | 31 | 43 | 33 | 52 |
| 100 | 27 | 30 | 104 | 105 | 13 | 55 | 39 | 46 | 32 | 57 |

Rat 3

| Bin Size (msec) | Day1 | Day2 | Day3 | Day4 | Day5 | Day6 | Day7 | Day8 | Day9 | Day10 | Day11 | Day12 | Day13 | Day14 | Day15 |
|---|---|---|---|---|---|---|---|---|---|---|---|---|---|---|---|
| 3 | 47 | 51 | 75 | 111 | 87 | 102 | 65 | 78 | 41 | 37 | 84 | 68 | 89 | 137 | 72 |
| 5 | 46 | 60 | 68 | 109 | 79 | 91 | 59 | 67 | 40 | 33 | 93 | 81 | 87 | 111 | 54 |
| 10 | 39 | 67 | 64 | 136 | 98 | 89 | 66 | 79 | 46 | 41 | 116 | 103 | 105 | 142 | 79 |
| 25 | 128 | 215 | 193 | 482 | 359 | 313 | 165 | 270 | 174 | 127 | 373 | 399 | 446 | 567 | 291 |
| 35 | 134 | 287 | 295 | 683 | 538 | 456 | 225 | 382 | 255 | 206 | 447 | 437 | 637 | 749 | 409 |
| 50 | 200 | 390 | 380 | 994 | 715 | 743 | 322 | 546 | 372 | 284 | 603 | 585 | 937 | 918 | 505 |
| 65 | 224 | 448 | 474 | 1242 | 922 | 996 | 404 | 733 | 489 | 430 | 774 | 609 | 970 | 1156 | 602 |
| 75 | 279 | 531 | 526 | 1445 | 1109 | 1247 | 396 | 799 | 586 | 482 | 908 | 740 | 983 | 1205 | 719 |
| 90 | 367 | 613 | 600 | 2116 | 1399 | 1496 | 483 | 1081 | 672 | 632 | 1245 | 955 | 1354 | 1474 | 270 |
| 100 | 440 | 728 | 751 | 2398 | 1670 | 1938 | 549 | 1263 | 882 | 723 | 1452 | 1144 | 1427 | 1581 | 907 |

Rat 3 – Continued

| Bin Size (msec) | Day16 | Day17 | Day18 | Day19 | Day20 |
|---|---|---|---|---|---|
| 3 | 81 | 94 | 56 | 72 | 49 |
| 5 | 83 | 77 | 62 | 68 | 38 |
| 10 | 121 | 106 | 58 | 72 | 41 |
| 25 | 375 | 376 | 218 | 206 | 186 |
| 35 | 458 | 633 | 360 | 320 | 264 |
| 50 | 532 | 857 | 480 | 477 | 356 |
| 65 | 617 | 979 | 503 | 514 | 419 |
| 75 | 734 | 1059 | 602 | 546 | 468 |
| 90 | 900 | 1308 | 746 | 687 | 595 |
| 100 | 895 | 1416 | 760 | 734 | 686 |

Rat 4

| Bin Size (msec) | Day1 | Day2 | Day3 | Day4 | Day5 | Day6 | Day7 | Day8 | Day9 | Day10 | Day11 | Day12 | Day13 | Day14 | Day15 |
|---|---|---|---|---|---|---|---|---|---|---|---|---|---|---|---|
| 3 | 2 | 1 | 17 | 13 | 10 | 6 | 25 | 9 | 4 | 8 | 5 | 1 | 5 | 4 | 4 |
| 5 | 3 | 1 | 20 | 11 | 13 | 4 | 30 | 10 | 3 | 10 | 4 | 4 | 6 | 7 | 4 |
| 10 | 9 | 3 | 26 | 17 | 22 | 3 | 24 | 13 | 3 | 5 | 8 | 4 | 11 | 5 | 8 |
| 25 | 18 | 11 | 56 | 51 | 50 | 11 | 71 | 27 | 8 | 10 | 20 | 15 | 16 | 9 | 25 |
| 35 | 15 | 11 | 70 | 49 | 43 | 8 | 92 | 37 | 9 | 19 | 23 | 12 | 16 | 18 | 27 |
| 50 | 13 | 9 | 87 | 51 | 59 | 8 | 105 | 45 | 11 | 17 | 24 | 14 | 14 | 18 | 31 |
| 65 | 15 | 12 | 86 | 62 | 68 | 7 | 99 | 39 | 15 | 20 | 30 | 11 | 16 | 16 | 34 |
| 75 | 19 | 9 | 92 | 64 | 67 | 5 | 110 | 48 | 14 | 20 | 24 | 11 | 9 | 20 | 36 |
| 90 | 21 | 10 | 90 | 71 | 89 | 6 | 96 | 42 | 13 | 21 | 35 | 12 | 11 | 19 | 41 |
| 100 | 17 | 14 | 91 | 64 | 83 | 7 | 104 | 43 | 15 | 26 | 38 | 15 | 10 | 20 | 40 |
